# Chaperone-assisted cryo-EM structure of *P. aeruginosa* PhuR reveals molecular basis for heme binding

**DOI:** 10.1101/2023.08.01.551527

**Authors:** Paweł P. Knejski, Satchal K. Erramilli, Anthony A. Kossiakoff

## Abstract

Pathogenic bacteria, such as *Pseudomonas aeruginosa*, depend on scavenging heme for the acquisition of iron, an essential nutrient. The TonB-dependent transporter (TBDT) PhuR is the major heme uptake protein in *P. aeruginosa* clinical isolates. However, a comprehensive understanding of heme recognition and TBDT transport mechanisms, especially PhuR, remains limited. In this study, we employed single-particle cryogenic electron microscopy (cryo-EM) and a phage display-generated synthetic antibody (sAB) as a fiducial marker to enable the determination of a high-resolution (2.5 Å) structure of PhuR with a bound heme. Notably, the structure reveals iron coordination by Y529 on a conserved extracellular loop, shedding light on the role of tyrosine in heme binding. Biochemical assays and negative-stain EM demonstrated that the sAB specifically targets the heme-bound state of PhuR. These findings provide insights into PhuR’s heme binding and offer a template for developing conformation-specific sABs against outer membrane proteins (OMPs) for structure-function investigations.

## INTRODUCTION

TonB-dependent transporters (TBDT) are integral outer membrane (OM) proteins found in Gram-negative bacteria. TBDTs play vital roles in the acquisition of essential nutrients that are either too large or scarce to be taken up by simple diffusion through the semi-permeable OM, including iron, heme, vitamin B12, and carbohydrates^1,2^. These OM transporters form transiently open channels that enable the selective uptake of substrates into the periplasmic space while preserving the integrity of the outer membrane^2,3^. TBDTs are physically coupled to an inner membrane energy transducing complex, which allosterically powers nutrient uptake into the periplasm using the proton motive force, the details of which are not completely understood^4–6^.

Iron is one such essential nutrient for the survival and virulence of many bacterial pathogens, and hence iron complexes, including heme, constitute the majority of substrates taken up by TBDTs^3,7,8^. *Pseudomonas aeruginosa* is an opportunistic pathogen with at least two heme uptake systems: the heme assimilation system (*has*) and the *Pseudomonas* heme uptake (*phu*) system^9,10^. The *has* operon encodes a secreted hemophore, HasA, which extracts heme from environmental hemoglobin and delivers it to an OM TBDT, HasR^11,12^. The *phu* operon encodes an 82-kDa OM TBDT, PhuR, that captures heme directly from the extracellular milieu^9,13^. Substrate binding to each TBDT triggers a signal transduction cascade across the membrane, with the TonB-ExbB-ExbD complex catalyzing heme uptake into the periplasmic space. The presence of multiple systems, common in Gram-negative bacteria, thus provide *P. aeruginosa* a significant advantage in competing for this limited resource within the host environment^10^. Moreover, iron acquisition is known to be essential for biofilm formation^13^.

It is in this setting that PhuR appears to play a crucial role. Transcriptome analyses from *P. aeruginosa* clinical lung isolates show PhuR is the major OM heme receptor in clinical infections, rather than HasR, with mutations in the *phu* operon resulting in increased PhuR expression levels^14^. *P. aeruginosa* Δ*phuR* knockout strains are unable to utilize heme as well as the wild-type, suggesting an important role for PhuR^9^. These observations suggest potential physiological roles for each system, with HasA/HasR thought to functional primarily in heme sensing while PhuR is mainly involved in uptake^13^. Biochemical analysis of PhuR suggest a His-Tyr pair is involved in heme binding, but limited structural evidence is currently available for heme recognition by TBDTs^10^.

To better understand the structural basis for heme recognition by PhuR, we endeavored to determine its three-dimensional structure using single particle cryogenic electron microscopy (cryo-EM). Due to the current size limitations of cryo-EM, we assisted our efforts by the generation of a synthetic antibody (sAB) specific for PhuR using a robust phage display technology developed in our group^15–18^. These sABs have demonstrated utility as fiducial markers in cryo-EM^19–22^. We determined a 2.5 Å resolution cryo-EM structure of PhuR using one sAB from this campaign, termed sAB11. Our structure contains bound heme and an LPS molecule, both co-purified with PhuR, providing structural insights into heme recognition and lipid interactions with OM proteins. In contrast with a previous biochemical study in which PhuR was identified as dependent on Y519 and H124 for heme binding, our structure reveals that Y529 plays a pivotal role in coordinating heme near the top of the extracellular domain^10^. We further demonstrate that sAB11 is conformation-specific by showing it can bind to holo-PhuR, but not apo-PhuR. Our results provide insight into substrate recognition by TBDTs and serve as a template for the generation of biochemical tools against similar OMPs.

## RESULTS

### Purification of *P. aeruginosa* PhuR and synthetic antibody generation

We recombinantly expressed *Pseudomonas aeruginosa* PhuR using an N-terminal hexa-histidine tag and by replacing the native signal peptide (spanning residues 1-25) with a PelB sequence^23^. PhuR was expressed in *E. coli* C41 (DE3) cells, solubilized from outer membranes using DDM, and purified by immobilized metal affinity chromatography (IMAC). DDM-solubilized PhuR was then analyzed by gel filtration chromatography, and eluted as a highly pure single peak from the column (**Supplemental Fig. 1A)**. Circular dichroism (CD) spectroscopy confirmed the purified detergent-solubilized protein was primarily comprised of β-sheets, and thermal denaturation experiments showed purified PhuR has a melting temperature (T_m_) of 71°C (**Supplemental Fig. 2**).

To overcome current cryo-EM size limitations, we screened a highly diverse and fully synthetic antibody (sAB) library to obtain binders specific for recombinant PhuR^24^. PhuR was reconstituted into lipid-filled nanodiscs (**Supplemental Fig. 1C**) formed using chemically biotinylated membrane scaffold protein (MSP) E3D1. The nanodisc provides PhuR a lipid environment to ensure the protein a natural environment during the phage display selection procedure. Five rounds of biopanning were performed using a well-established selection protocol^17,18,25^ increasing stringency round to round by systematically lowering protein concentration from 200 nM in round 1 to 25 nM in round 5. Notably, no heme was added to the selection buffer. We screened 96 clones from the final round phage pool for the initial validation and, through sequencing, identified over 70 unique PhuR-specific binders. We then selected the 18 candidates with highest binding signal from this pool for subcloning, expression, purification, and further analysis (**Fig. 1A-B, Supplemental Table 1**). This set was then filtered based on sAB affinities for PhuR and complex formation efficiency (**Fig. 1C**, **Supplemental Table 2**). Based on this EC_50_ analysis, sAB11, which has an EC_50_ of ∼9 nM, was chosen for structure determination of PhuR by fiducial-assisted cryo-EM.

**Figure 1:**
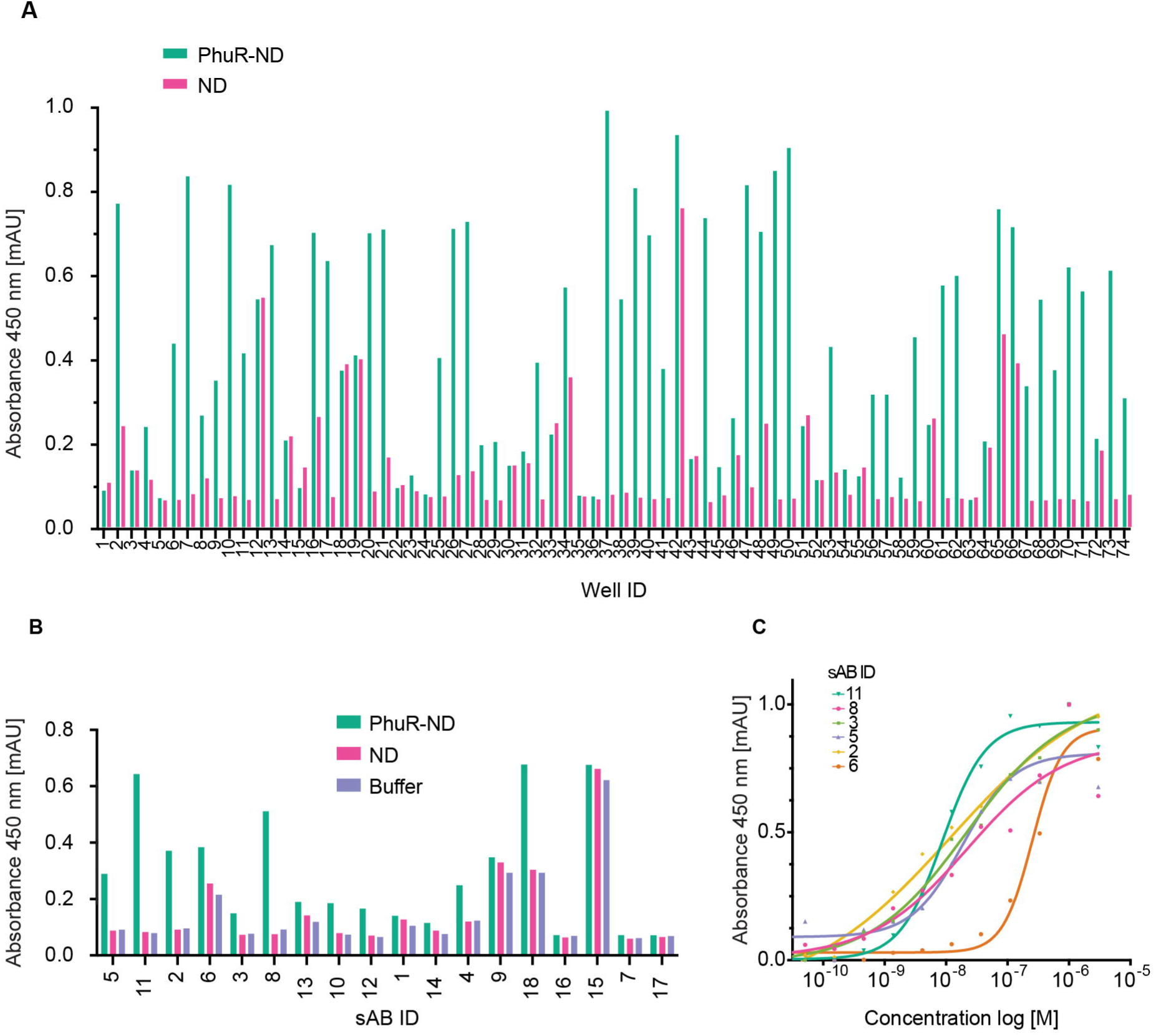
sAB generation and validation **A.** Single point phage ELISA comparing binding of unique clones to PhuR-containing nanodiscs and empty nanodiscs as a control. **B.** Single-point ELISA signal of purified selected sAB clones from the initial phage pool (panel A). Binding of 300 nM of purified sABs tested using PhuR-containing nanodiscs (green), empty nanodiscs (raspberry pink), and a buffer only control (purple). **C.** Affinity estimation ELISA for select sABs measuring binding to PhuR-containing nanodiscs. Data are background subtracted and normalized. Plots are fitted using a nonlinear regression model with variable slope in Prism 9.0 to determine EC_50_.

### sAB-assisted single-particle cryo-EM enables high-resolution structure determination of PhuR

For cryo-EM single-particle analysis (SPA), PhuR was reconstituted into amphipols, which have demonstrated utility in cryo-EM SPA of membrane proteins^26^. Amphipol-reconstituted PhuR similarly forms a stable complex with sAB11 as PhuR reconstituted into nanodiscs (**Supplemental Fig. 1C, E-F**). To improve the cryo-EM data, we also utilized a nanobody (Nb) that binds with high-affinity to the sAB hinge region and functions as an additional fiducial marker for cryo-EM SPA^27^. We obtained high-quality grids of the tripartite complex and collected cryo-EM data (**Supplemental Fig. 3A**).

Using this sample preparation, we determined the structure of the PhuR-sAB11-Nb complex to an overall nominal resolution of 2.5 Å, with most of the PhuR structure resolved to ∼2.4 Å (**Fig. 2A**, **Supplemental Fig. 3A-C**). The majority of the cryo-EM data contained monomeric PhuR; however, it also included an antiparallel dimer form of PhuR, which could be readily segregated into a separate particle class. The two distinct particle classes are apparent from two-dimensional class averages, and the intact complexes of PhuR with sAB11 and Nb are clearly visible (**Supplemental Fig. 3A**). The antiparallel PhuR dimer similarly contained two molecules of the sAB11-Nb complex and had a nominal 3 Å resolution. Given the non-physiological nature of the dimer and overall lower resolution, it was not further analyzed.

**Figure 2:**
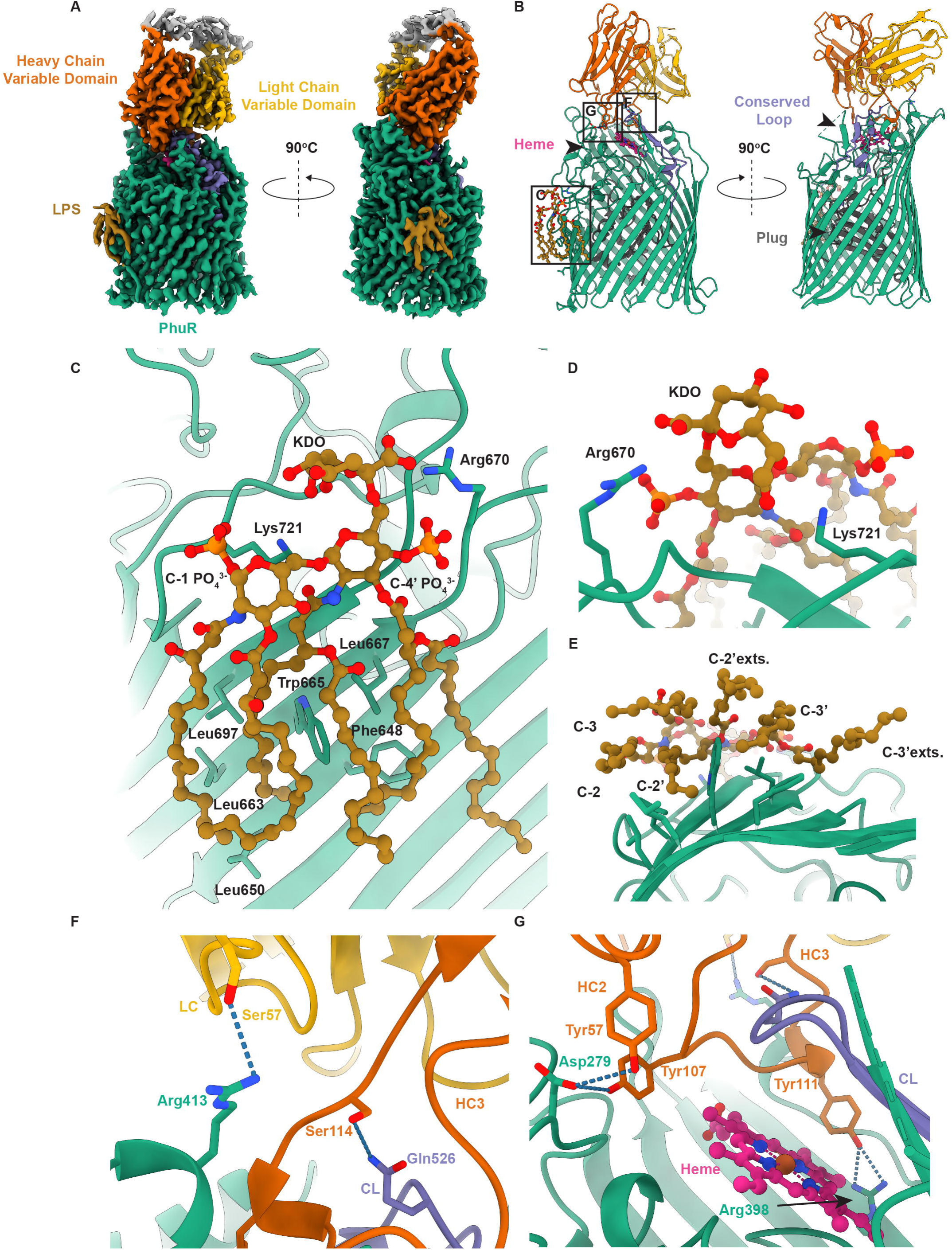
CryoEM structure of PhuR-sAB11 complex **A.** Cryo-EM density map of PhuR (green) bound to sAB11 (heavy chain (HC) shown in orange and light chain (LC) in golden brown) and LPS (mustard). Regions excluded from the model are white. **B.** Front and side views of PhuR-sAB11 complex with LPS and bound heme (raspberry pink) shown. The complex is colored as in A. The conserved loop (CL) with heme-coordinating Y529 colored purple. **C.** Side view of LPS and PhuR residues potentially interacting with it. **D.** Top view of LPS KDO sugar and Lipid A headgroup showing conserved interactions with R670 and K721. **E.** Bottom view of LPS tails showing hydrophobic interactions with PhuR. **F.** Close-up view of LC interactions with PhuR. **G.** Close-up view of HC complementarity-determining regions (CDRs) 2 and 3 interactions with PhuR.

For initial cryo-EM density fitting, we started with a model of monomeric PhuR comprising residues 47-578 and 585-764 (**Table 1, Supplemental Fig. 4-5)** using AlphaFold^28^. The final experimental structure shows PhuR is an 22-stranded β-barrel outer membrane protein (OMP). The strands are all uniformly tilted approximately 45° to the normal of the membrane (as depicted in **Fig. 2B**) and the structure features an N-terminal plug domain situated within the barrel. These structural features are conserved amongst TonB-dependent transporters (TBDT)^2,3^. PhuR is overall asymmetric in shape, with strands 6-9 forming a 74 Å high wall along one side of the barrel that extends well into the extracellular domain, and shorter wall on the other side of the barrel (**Fig. 2B**). We identified density along the shorter outer wall of the barrel, near strands 16-20, that was tentatively assigned as a portion of a lipopolysaccharide (LPS) molecule that was present during purification (**Fig. 2A-B**). We could resolve the Lipid A and first KDO sugar in this density, while the rest of the LPS was not visible in the cryo-EM map (**Fig. 2C-D**). The structure revealed the Lipid A C-1 and C-4’ phosphate headgroups likely interacting with PhuR residues K721 and R670, respectively, while a number of conserved hydrophobic residues appear to form van der Waals interactions with the six fatty acid tails (**Fig. 2E**). Headgroup interactions involving both arginines and lysines are commonly observed features in LPS binding sites, with recent evidence of such, including hydrophobic residues with the acyl chains, demonstrated in the case of the ferrichrome transporter FhuA (PDB ID 8A8C)^29^. Another study, based on molecular dynamics simulations, suggests that the proximity of arginine residues to the Lipid A phosphate groups supports the interpretation that these residues play pivotal roles in LPS binding^21^.

Residues 47-166 form the N-terminal plug domain, a hallmark feature of TBDTs that is critical for sensing and uptake of the ligand^3,30^. The N-terminal residues preceding the plug, which exclude the processed signal peptide but potentially include the Ton box, were not resolved^2^. These residues are resolved in some structures, including BtuB (PDB ID 1NQH), but has also been shown to be more dynamic when substrates are bound in other TBDTs^6,31,32^. The rest of the plug was well resolved and contained the conserved fold typical of β-barrel TBDTs^3^. The core of this domain is a 4-stranded β-sheet tilted with respect to the plane of the membrane, while the remainder is composed of short helices and long loops projecting upwards towards the extracellular surface (**Fig. 2B**).

The periplasmic loops, or turns as they are conventionally described, of PhuR are all relatively short. The longer of the extracellular loops, many composed of well-structured elements, collectively form a vestibule (∼1780 Å^3^ volume) above the hydrophilic interior of the barrel, which is 40 Å wide and occluded by the plug in our structure. We could confidently model all of these segments except for the tip of the L8 connecting strands 15 and 16 (residues 579-584), which appears to be disordered in the experimental structure. **(Supplemental Fig. 4A-B).**

The majority of the loops in the experimental PhuR structure have identical or similar conformations to the AlphaFold model, with RMSD values of 0.7-2.0 Å for nine of the eleven loops. The two notable exceptions were the long extracellular loops 10 (residues 669-694) and 11 (718-754), which, when compared to the AlphaFold model, have backbone RMSD values of 3.9 and 4.8 Å, respectively. These loops, particularly L11, converge towards the center of the extracellular section of the barrel, effectively creating a seal that encloses the vestibule **(Supplemental Fig. 4C).**

The sAB11 interacts extensively with the extracellular vestibule of PhuR, primarily through complementarity-determining regions (CDRs) H2, H3, and L3. The contacts are formed by both hydrogen bond and van der Waals interactions with the extracellular loops L3, L4, L5, and L7, and bury ∼1000 Å^2^ of surface area at the interface (**Fig. 2F-G**). The 16-residue long sAB CDR-H3 forms a blunt wedge, packing against an extracellular conserved loop (CL). This loop, L7, contains the conserved FRAP/PNPNL motif that spans residues 510-513 and 533-537, respectively and the β-strands that comprise the taller edge of the barrel (**Fig. 2B, Supplemental Fig. 4B**). This conserved motif, which is found in all TBDTs, which transport heme and is the loop that contains the axial heme ligand, diverges somewhat in PhuR, where the sequence is actually FRTP^33,34^. This is the loop that contains Y529, the axial ligand in PhuR. Hydrogen bond interactions mediate many of the contacts between sAB11 and PhuR, including CDR-H3 residue S114 with CL residue Q526, HC residues Y57 and Y107 with PhuR D279, and H3 residue Y111 with R398. The latter two PhuR residues are located on the tall edge of the barrel within extracellular loops and β-strands, respectively. Notably, the conformations of the extracellular loops forming the sAB epitope are identical or similar to the conformations predicted in the AlphaFold model.

### PhuR cryo-EM map revealed heme coordinated by conserved loop residue Y529

We observed well-resolved non-protein cryo-EM density in the extracellular vestibule near the sAB epitope, which we readily modeled as a bound heme molecule that co-purified with PhuR (**Supplemental Fig. 5**). The presence of heme was readily apparent by visual inspection of purified PhuR and confirmed by analysis of the UV-vis spectra of the PhuR-sAB11-nanobody complex (**Supplemental Fig. 1D, Fig. 2G**, **Fig. 3A-C**, **Fig. 5A**). Additionally, the fact that both the antiparallel dimers and monomer classes of PhuR, which constitute 85% of the particles present after several rounds of 2D classification, displayed identical densities strongly implies that the heme-bound form of PhuR was predominant in solution (**Supplemental Fig. 3A**).

**Figure 3:**
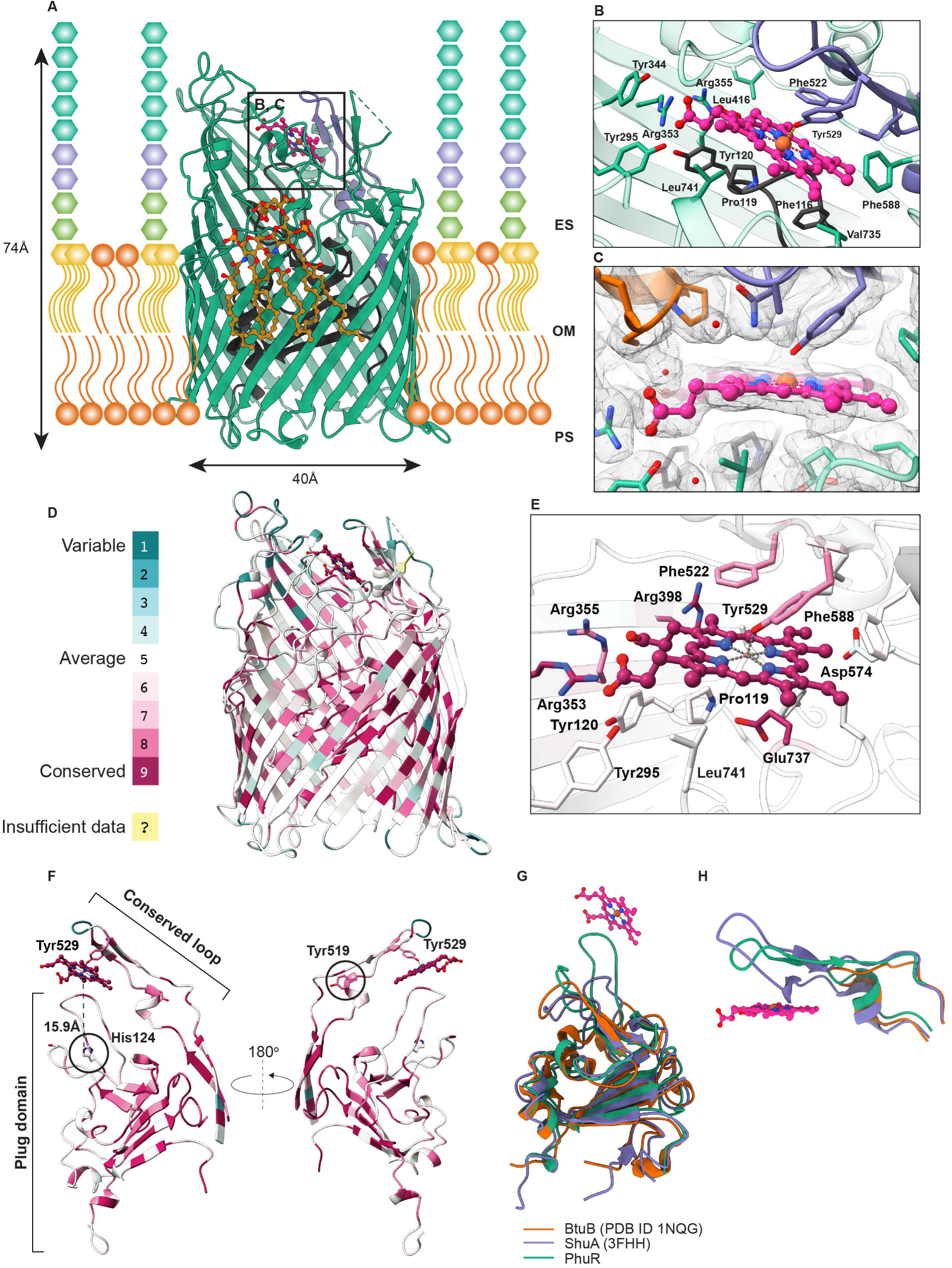
Heme recognition by PhuR **A.** Side view of PhuR placed in a schematic lipid bilayer with LPS molecules with indicated Lipid A (yellow), inner core sugars (green), outer core sugars (purple), O-antigen (green). The plug domain (dark gray), bound LPS, conserved loop, and heme are also visualized. PhuR dimensions in Å indicated with arrows. ES - extracellular space, OM - outer membrane, PS - periplasmic space. **B.** Residues involved in the heme (raspberry pink) interaction, including the coordinating Y529. Plug domain and conserved extracellular loop colored dark gray and purple, respectively. **C.** Heme binding site overlaid with cryo-EM density map (gray mesh) contoured to visualize the model and proximal water molecules. **D.** PhuR cryo-EM structure colored according to evolutionary conservation scores of individual residues^45,46^. Legend for conservation scores shown below structure. **E.** Close-up view of heme binding site to show conservation of interacting residues. **F.** Conservation of heme-binding extracellular loop and plug domain shown with bound heme. **G.** Overlay of plug domain structures of PhuR (this study), ShuA (3FHH)^31,36^, and BtuB (1NQG)^31^ to compare conformations of the heme binding loop. Heme molecule in PhuR structure shown and colored as above. **H.** Overlay of conserved extracellular loops from the same three structures to highlight conformational differences.

The heme molecule is bound through multiple sets of interactions with the protein. The heme iron is coordinated through the hydroxyl group of Y529, which is located on the structured extracellular CL between strands 13-14 that forms a β-turn and contacts sAB11 CDR-H2. The distance between the Y529 hydroxyl oxygen and heme iron (1.9 Å) is consistent with the typical Fe-O distance observed in heme-coordinating tyrosine residues^35^. The aforementioned conserved FRTP (FRAP) and PNPNL motifs flank the conserved Y529, with the CL forming a lid over the extracellular heme-binding site to create a predominantly hydrophobic binding pocket near the outermost surface of PhuR (**Fig. 3B, D-F, Supplemental Fig. 6**). Several nearby residues stabilize heme in this pocket, including R353 and R355, which form hydrogen bonds with the propionate sidechains, F522 and F588 through *π*-*π* stacking, and L416, I531, V735, and L741 through van der Waals interactions (**Fig 3B, E, Supplemental Fig. 7**). At the tip of a 24-amino acid long loop rising upwards from the plug domain are residues F116, P119, and Y120, which also interact with heme, thus connecting the extracellular binding site with the region of PhuR responsible for partnering with inner membrane energy-transducing complexes to facilitate ligand uptake. While the conformation of the plug domain is similar to those observed in β-barrel OMP structures in their apo states, such as BtuB (PDB ID 1NQG) and ShuA (PDB ID 3FHH), the orientation and substrate interactions of this loop are distinct in our PhuR structure **(Fig. 3G-H**)^31,36^. Combined with the lid formed by the CL, this plug loop sandwiches the heme into its binding site. This is structurally similar to the HasA-HasR complex (PDB ID 3CSL), where the heme iron is coordinated after transfer from HasA by HasR to residues H603, which resides in the same extracellular loop, and H189 from the plug loop^12^. In the BtuB structure with bound cyanocobalamin (1NQH), the plug, including the homologous loop, interacts extensively with the substrate, in part due to the binding site residing at a lower position within the barrel and due to the substrate size^31^.

Biochemical and mutational analyses previously reported that the conserved residues Y519, located on the CL following the FRAP motif, and H124, on the plug domain, are involved in heme coordination^10^. In the structure reported here, Y519 is pointing away from heme, with the hydroxyl positioned towards the extracellular space. H124, which is part of the same plug domain loop that contacts the heme, is located too far, ∼16 Å, to participate in heme binding (**Fig. 3D-F**), and is instead likely forming hydrogen bond interactions with plug residues E85 or Q122. Neither H124 nor Y519 are located near the vestibule where heme is observed to be bound (**Fig. 4A**). Notably, the conformations of H124, Y519 and Y529 are essentially identical (0.74 Å RMSD) in the AlphaFold model. A second, much smaller cavity (536 Å^3^) spans the space between Y519 and H124 located below the observed heme binding site (**Fig. 4A-B**), with a narrow channel connecting the two cavities (**Fig. 4C-E)**. There was no cryo-EM density in the second cavity corresponding to heme; while it is apparently large enough in volume to accommodate heme (∼460 Å^3^), there are no potential ligands to accommodate its occupancy. The proposed Y519-H124 coordination would require conformational changes to correctly position these residues in this potential binding pocket. Moreover, access to this pocket for heme would require an expansion of the extracellular region of PhuR, as the pathway to the Y519-H124 from the vestibule is constricted in the structure by L10 (**Fig. 4E**). Therefore, if this cavity represents a second ligand binding site lower in the barrel along the uptake pathway, as has been proposed for homologous TBDTs from other studies, a conformation of PhuR distinct from both the experimental structure and AlphaFold model would be required to enable heme binding at this site and coordination by Y519 and H124^37–39^.

**Figure 4:**
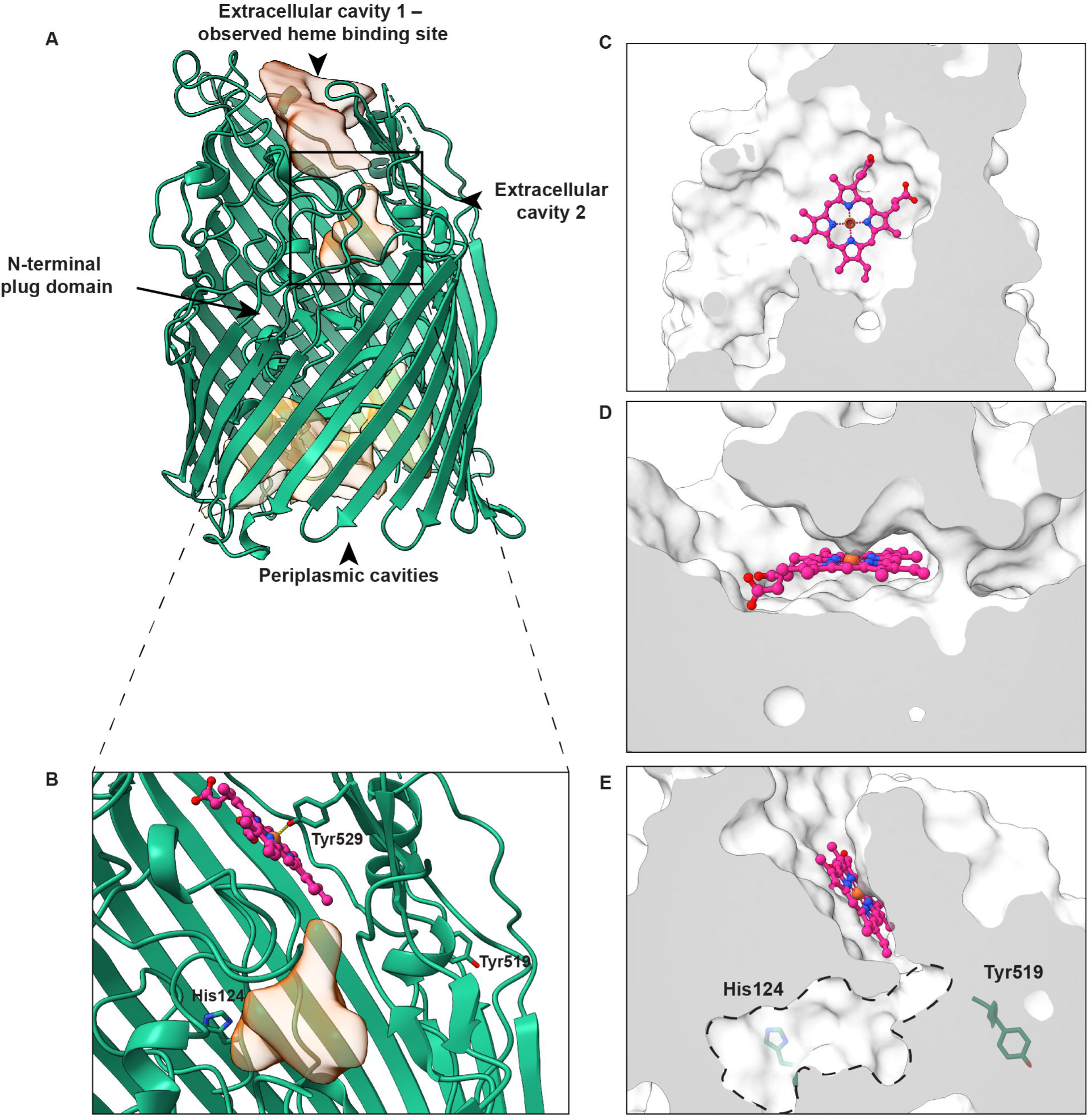
Visualization of PhuR cavities suggest a potential heme uptake mechanism. **A..** Side view of PhuR protein in cartoon representation with extracellular and periplasmic cavitites shown as transparent volumes. Heme is bound to PhuR in the extracellular cavity 1. The second extracellular cavity is proposed based on biochemical data from a previous study^10^. **B.** Close-up view of extracellular cavity 2 with H124 and Y519 shown as sticks together with the bound heme and coordinating Y529. **C.** Top and **D.** side views of capped PhuR structure in surface representation with heme in magenta. **E.** Side view of second cavity bounded by dashed lines with His124 and Tyr519 represented as sticks showing a narrow channel connecting the two cavities.

**Figure 5:**
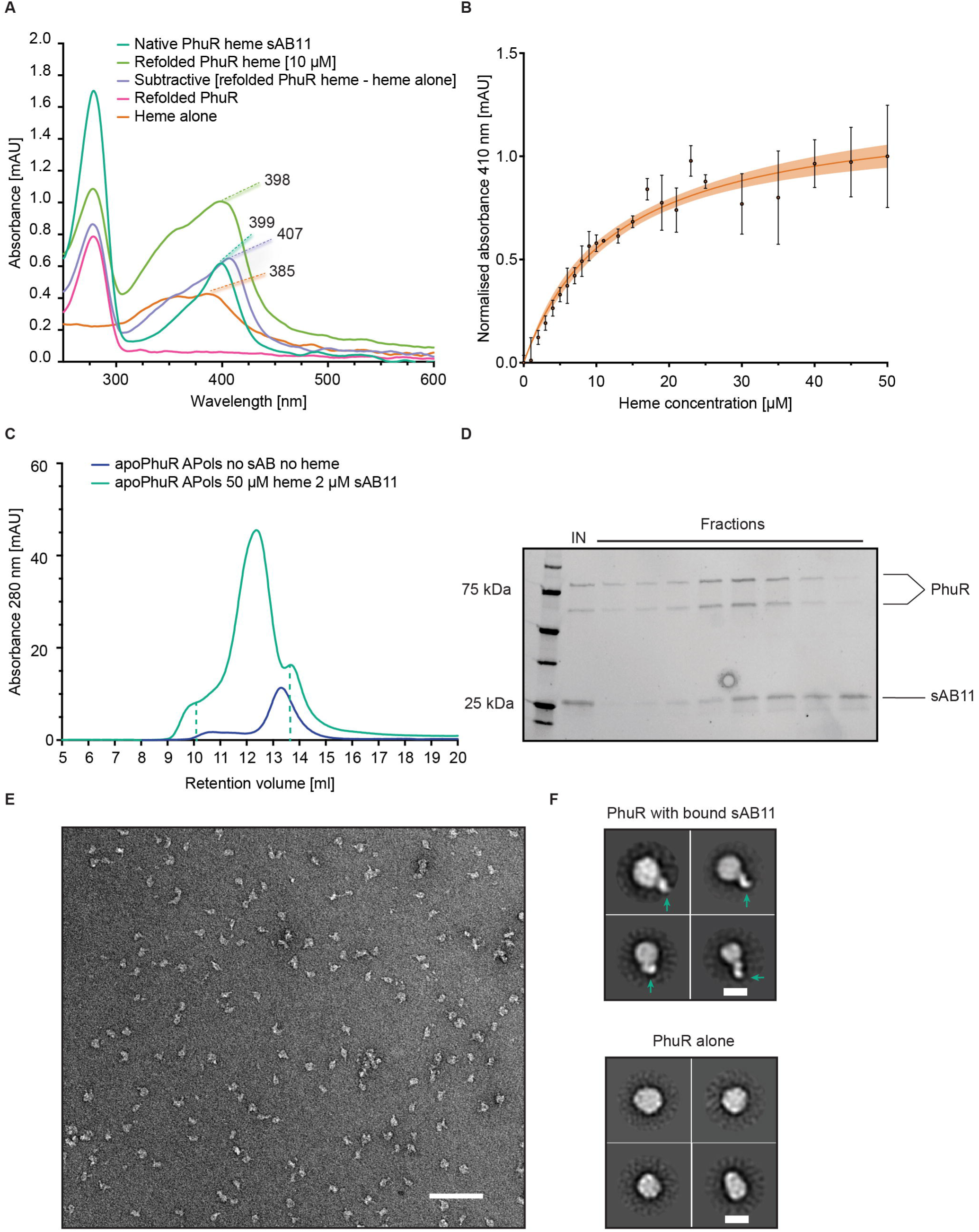
Biochemical analysis of refolded PhuR and its interaction with sAB11**. A.** UV-Vis spectra for native sAB-bound PhuR, heme alone, refolded PhuR in the presence and absence of heme, and the spectrum resulting from the subtraction of heme alone from refolded PhuR with addition of heme. **B.** Spectroscopic evaluation of heme binding to refolded PhuR using increasing concentrations of heme, performed in triplicate. Data are plotted using Prism (v9.0) software and represented as mean ± SEM, with the 95% confidence intervals shown (orange). **C.** Overlay of chromatograms from size exclusion column (SEC) for refolded PhuR alone (blue) and after incubation with heme and sAB11 (green). Dashed lines indicate collected fractions. **D.** SDS-PAGE gel of elution collected after SEC from panel C of refolded PhuR in the presence of heme and sAB11. PhuR is present as two bands due to the heat modifiability property of OM proteins^47^. **E.** Representative ns-EM micrograph of peak fraction from panel C. Scale bar represents 50 nm. **E.** 2D class averages of refolded PhuR with sAB11 (green arrows) and heme (top panel) and PhuR alone (bottom panel). Both samples are reconstituted into amphipols. Scale bar represents 10 nm.

### Refolded PhuR is active for heme binding

To better understand the conformational changes of PhuR associated with heme binding, we endeavored to isolate the apo state by expressing into inclusion bodies PhuR without the signal peptide. This method is well-established for obtaining high yields of β- barrel outer membrane proteins for structural studies^23^. PhuR was denatured using 5 M urea, applied to an IMAC column, refolded by slow buffer exchange in the presence of DDM, and subsequently purified. Gel filtration chromatography was used to separate folded and unfolded proteins. Peak fractions containing folded PhuR were pooled and analyzed using CD spectroscopy, which established that refolded PhuR is composed predominantly of β-sheet secondary structure (**Supplemental Fig. 2**). Thermal denaturation of refolded PhuR showed a T_m_ of 65°C, 6°C lower than the natively expressed version. Spectroscopic analysis of refolded PhuR suggested heme was not present in the sample (**Fig. 5A**). Employing a spectrophotometric assay, we determined that refolded PhuR binds hemin, eliciting a redshifted spectrum in the Soret region compared with unbound heme. The observed shift is characteristic of the typical spectral changes associated with both unbound and bound heme, and the wavelength of the Soret peak was nearly identical in the natively expressed PhuR-sAB11 complex (**Fig. 5A**)^9,10^. We observe that, employing this assay, the estimated affinity for the binding of hemin to refolded PhuR is approximately 12.5 µM (**Fig. 5B**). This affinity appears lower than anticipated, given the consistent presence of an intact heme-loaded complex throughout multiple rounds of washing steps during the phage display biopanning process for native PhuR. We speculate that the compromised affinity of this heme analog may stem from two potential factors: its low solubility and the likelihood that the refolded material could encompass a population of non-natively folded molecules.

The ∼70 sABs obtained from the native PhuR selection were tested for binding to refolded PhuR. None of these sABs showed measurable binding to the refolded protein, including sAB11. This was not surprising since in the absence of heme, which was not observed using spectrophotometry, and LPS, which is not bound to proteins refolded from inclusion bodies, parts of PhuR are likely highly flexible and dynamic. Relatedly, in several crystal structures of β-barrel OMPs, density for extended loops is often missing due to flexibility, and it is known these regions are essential for transport^31,36,40–42^

To better understand the mode of sAB binding, we formed a complex between sAB11 and refolded PhuR in the presence of hemin. Amphipol-reconstituted refolded PhuR was incubated with 50 µM hemin, and subsequently sAB11 was added to this solution. The complex was purified by gel filtration chromatography (**Fig. 5C**). Analysis of the peak fractions by SDS-PAGE showed bands for both PhuR and sAB11 when hemin was present (**Fig. 5D**). The PhuR-sAB11 complex was not formed when hemin was not present. This complex was subsequently analyzed by negative-stain electron microscopy (ns-EM). Two-dimensional ns-EM class averages clearly showed the sAB bound to refolded PhuR in a manner similar to native PhuR (**Fig. 5E-F**). These experiments demonstrated both that sAB11 is specific to the heme bound conformation of PhuR, and that the mode of sAB binding could be recapitulated by the addition of hemin to the refolded PhuR preparation.

## DISCUSSION

Iron is an essential nutrient for all bacteria, and establishing a ready source is indispensable for growth and virulence for a number of pathogens^3,7^. PhuR is a central player in heme acquisition in *Pseudomonas aeruginos*a, and is particularly important in clinical strains of this nosocomial pathogen^9,10,13^. It is thus potentially an attractive target for therapeutics, and understanding its structure-function relationship is key for the design and development of drugs against this TonB-dependent transporter.

We describe here the first experimental structure of PhuR using single particle cryo-EM. To overcome the current size limitation of cryo-EM for this 82-kDa integral membrane protein, we generated a sAB11, using a phage display technology developed in our group for nanodisc-reconstituted membrane proteins^15–18^. High-quality cryo-EM data was obtained for the complex of PhuR with sAB11, which facilitated accurate model building and detailed interpretation at 2.5 Å resolution.

The overall architecture is consistent with other TBDTs, with PhuR composed of 22 antiparallel β-strands and an N-terminal plug domain. Long extracellular loops form a large vestibule, which contained a bound heme molecule. The cryo-EM density contained well resolved features, enabling model building with high accuracy the interactions facilitating heme binding and recognition with high accuracy. Notably, the heme molecule was co-purified with PhuR and remained bound through multiple washing steps during the phage display selections suggesting it associates with very high affinity to the native protein. The conformation of the sAB epitope, composed of several extracellular loops, in both the cryo-EM structure and the AlphaFold model were remarkably similar. The major differences were in extracellular loops L10 and L11, which completed the heme binding site. The heme is coordinated by a Fe-O bond formed between the heme iron and the PhuR Y529 phenoxyl. This observation adds to a growing body of knowledge regarding the role of tyrosine in heme binding for a variety of proteins^10,13,43^. The location of the observed heme binding site contrasts with earlier biochemical and mutational analyses suggesting pivotal roles for Y519 and H124^10^. These residues are distal to the observed heme binding site, but instead identified proximal to a secondary, smaller cavity situated lower in the barrel and closer to the plug domain. No discernible class averages in the data indicated additional minor substates, leaving unanswered whether this second cavity might represent a potential pathway to an intermediate state in heme transport. The concept of two-site binding, previously inferred for PfeA from *P. aeruginosa* and *E. coli* FepA, aligns with the presence of at least two binding sites for substrates in those TBDTs^37–39^. However, both our experimental structure and the AlphaModel model indicate that a significant structural reconfiguration in both the extracellular loops and plug domain, perhaps induced by TonB binding, are necessary for enabling heme binding at this potential second site.

To better understand the conformational changes associated with heme binding, we denatured and refolded PhuR to obtain its apo state, and determined its ability to bind hemin. In the hemin bound form, refolded PhuR was able to form a complex with sAB11, demonstrating the conformation-specificity of this reagent. Negative-stain EM analysis confirmed the binding of sAB11, showing we could recapitulate its binding mode using the refolding preparation together with added hemin. Considering the heme-bound form was utilized for sAB biopanning and the close proximity of the sAB epitope to the heme-binding site, the specificity of the sAB for this particular conformational state is entirely expected. Our group has previously shown that a necessary prerequisite to obtaining conformation-specific sABs is the stabilization of the target protein in the desired state^25,44^. In this case, the visual inspection, spectral analyses, and structural data all demonstrated that the heme bound form is highly enriched in the preparation of recombinant PhuR from *E. coli* membranes.

Overall, our findings shed light on the structural features of PhuR and its interactions with heme, adding to the growing body of knowledge of how TBDTs initially bind and recognize their transported substrates. Future studies could further investigate the functional relevance of heme binding to the observed site, the role of the second cavity, and the relevance of both to the transport mechanism of PhuR. Alternative strategies could enable the structure determination of additional functionally-relevant states of PhuR. These could include mutational scanning to identify residues in the vestibule and the plug domain along the transport pathway. Finally, our study underscores the effectiveness of our biopanning methodologies in generating highly specific binders against TBDTs.

## ACKNOWLEDGEMENTS

We would like to thank Khuram Ashraf, Tomasz Gawda, Duy P. Hua, Marc André Leibundgut, Julia Marcińska, Somnath Mukherjee, Nicholas Noinaj, Zachary P. Schaefer, and Michał Śmiga for their critical feedback, helpful discussions, and kind assistance during the course of this study and in the preparation of this manuscript. We thank Jotham Austin II, Tera Lavoie, James R. Fuller, Yimei Chen, and the rest of the staff at the University of Chicago Advanced Electron Microscopy facility for help with cryo-EM data collection. We thank Elena Solomaha and the University of Chicago Biophysics core for their help with CD spectroscopic experiments. This work was supported by NIH grant R01GM117372 to A.A.K.

## CRediT AUTHOR CONTRIBUTION STATEMENT

**Paweł P. Knejski** - Methodology, Investigation, Validation, Formal analysis, Visualization, Writing – original draft, Writing – review & editing. **Satchal K. Erramilli** – Supervision, Conceptualization, Methodology, Investigation, Validation, Formal analysis, Writing – original draft, Writing – review & editing, Project administration. **Anthony A. Kossiakoff** – Supervision, Conceptualization, Methodology, Validation, Resources, Writing – original draft, Writing – review & editing, Project Administration, Funding acquisition.

## STAR ★ METHODS RESOURCE AVAILABILITY

### Lead contact

Further information and requests for resources and reagents should be directed to and will be fulfilled by the Lead Contact, Anthony A. Kossiakoff.

### Materials availability

Primary data and other materials are available upon reasonable request to the lead contact.

### Data and code availability

The raw movie frames have been deposited into the Electron Microscopy Public Image Archive (EMPIAR), with accession code EMPIAR-11627. The cryo-EM density maps have been deposited into the Electron Microscopy Data Bank (EMDB, accession code EMD-41255). The coordinate file has been deposited in the Protein Data Bank (PDB, accession code 8THE). Synthetic antibodies generated in this study will be made available upon request. This paper does not report original code.

## EXPERIMENTAL MODEL AND STUDY PARTICIPANT DETAILS

The *E. coli* bacterial strains used for expression in this study are listed in the key resources table and growth conditions are described in detail in the Method Details section. Helper phage M13-KO7 was used for phage propagation in E. coli XL1-Blue cells.

## METHOD DETAILS

### Expression and purification of PhuR from *E. coli* outer membranes

The codon-optimized *phuR* gene from *Pseudomonas aeruginosa* PAO1 strain lacking the native signal sequence (MPLSPPFALRPCLALLLLSPSLALA) was synthesized and cloned into the expression vector pET20b(+), which contains an N-terminal pelB signal sequence and hexahistidine tag (Genscript). The construct was transformed into chemically competent C41 (DE3) cells (Lucigen) and plated after heat shock and recovery on LB agar supplemented with ampicillin. Colonies were allowed to grow with an overnight incubation at 37°C. The following day, a single colony was used to inoculate 30-mL of LB liquid media with 100 µg/mL ampicillin and grown overnight with shaking. The inoculum was transferred to 3 L of TB media supplemented with ampicillin and 0.4% (w/v) glycerol in baffled Fernbach flasks (Corning). Cultures were grown to mid-log phase (OD_600_ of 0.6-0.8), subsequently induced with 100 µM IPTG (RPI Chemicals), and proteins were expressed for 4 hours at 37°C. Cells were harvested by centrifugation and frozen until further use.

For purification, cell pellets were resuspended in lysis buffer (25 mM Tris, pH 7.4, 150 mM NaCl with 1 mM PMSF (Sigma) and homogenized (IKA Labs). Cells were lysed by two cycles of sonication (Branson Sonifier; 40% amplitude, 1.5-min cycle of 1-s on/1-s off). Membranes were solubilized by the addition of 1% n-dodecyl-β-D-maltopyranoside (DDM, Anatrace) followed by a one-hour incubation at 4°C with nutation. Insoluble material and cellular debris were removed by ultracentrifugation at 40,000 RPM for 45 minutes at 4°C using a Ti45 fixed-angle rotor (Beckman-Coulter). The resulting supernatant was supplemented with NaCl and imidazole to final concentrations of 500 and 20 mM, respectively, and subsequently applied to a 5-mL HisTrap column (GE Healthcare) pre-equilibrated with wash buffer (25 mM Tris, pH 7.4, 500 mM NaCl, 20 mM imidazole, 0.1% DDM). Following batch binding for one hour, the column was washed using a linear imidazole gradient (20-500 mM) formed by mixing wash buffer with elution buffer (wash buffer with 500 mM imidazole) on a fast protein liquid chromatography system (FPLC, ÄKTA) with fractionation and monitored using A_280_. Fractions containing protein were analyzed by SDS-PAGE, and PhuR-containing fractions were pooled and concentrated using 50-kDa MWCO centrifugal filters (Amicon). The resulting sample was dialyzed overnight in 25 mM Tris, pH 7.4, 150 mM NaCl buffer. The dialyzed sample was subsequently purified by size exclusion chromatography using a Superdex200 HiLoad 16/600 column (GE Healthcare). Fractions were analyzed by SDS-PAGE gel, and PhuR-containing fractions were pooled, concentrated, supplemented with 10% glycerol, flash frozen in liquid nitrogen, and stored at -80°C until further use. All protein concentrations were determined using the theoretical extinction coefficients.

### Expression and purification of PhuR from inclusion bodies

The codon-optimized *phuR* gene described above was also cloned into vector pET28b(+) (Genscript), which contains an N-terminal hexahistidine tag but lacks a signal sequence. A similar transformation and expression protocol was used, except using kanamycin (50 µg/mL) instead of ampicillin. Cells were harvested, homogenized, and lysed as described above. Inclusion bodies and other insoluble material were collected by centrifugation at 10,000×g at 4°C for 10 minutes. The resulting pellet was washed twice with denaturing buffer (25 mM Tris, pH 7.4, 500 mM NaCl, and 5 M Urea) and incubated for one hour at RT with nutation. The suspension was centrifuged at 10,000×g at 4°C for 20 minutes. The supernatant was collected, passed through at 0.2 µm filter, and subsequently injected onto a 5-mL HisTrap column (GE Healthcare) connected to an ÄKTA FPLC system. Prior to injection DDM was added to the supernatant at a final concentration of 0.1% (w/v). PhuR was refolded by slow exchange of buffer to remove urea using refolding buffer (25 mM Tris, pH 7.4, 500 mM NaCl, 0.1% DDM). Following refolding, PhuR was purified using a linear gradient of imidazole as described above for native PhuR. Fractions were analyzed by SDS-PAGE, and those containing PhuR were pooled, concentrated, and injected onto a Superdex200 HiLoad 16/600 SEC column (GE Healthcare) to separate folded and unfolded proteins. The SEC was performed using SEC buffer: 25 mM Tris, pH 7.4, 150 mM NaCl, and 0.02% DDM. Fractions were analyzed by SDS-PAGE using the heat modifiability property to assess whether PhuR was folded or unfolded^47^. Fractions containing folded PhuR pooled, concentrated, and stored as described above until further use.

### Circular dichroism and thermal denaturation assays

Far UV CD spectra were measured in a wavelength range of 180-260 nm using a Jasco J-1500 CD Spectrometer with a cell path length of 1 cm and a protein concentration of 5 µM in SEC buffer. The spectra were recorded using triplicate scans. For thermal denaturation studies, a fixed wavelength of 222 nM was monitored and simultaneous OD and CD scans were measured from 26 to 98°C. Temperature was increased by steps of 2°C with 3-min incubation periods between spectral measurements. Data fitting and analysis were performed using Spectra Manager (Jasco) and Prism 9.0 (GraphPad).

### Nanodisc reconstitution

MSP E3D1 was expressed and purified as previously described, including removal of the polyhistidine tag by digestion with TEV protease (kind gift of S. Koide)^48,49^. For phage display purposes, MSP was chemically biotinylated as previously described^17^. DDM-solubilized PhuR was mixed with either biotinylated or non-biotinylated E3D1 and total *E. coli* lipid extract (Avanti) in a 1:6:480 molar ratio. A lipid solution was prepared as previously described prior to incubation with the reconstitution mixture^50^. Empty nanodiscs for phage display were also prepared by mixing either biotinylated or non-biotinylated E3D1 with lipids in a 1:130 molar ratio. To form nanodiscs, detergent was removed by overnight incubation with activated Bio-Beads SM-2 (Bio-Rad). For purification of PhuR-containing nanodiscs, the reconstitution mixture was transferred from Bio-Beads to Ni-NTA resin (Qiagen). IMAC was performed using buffers similar to those described above, except without detergent. Fractions containing PhuR nanodiscs were pooled, concentrated, and injected onto a Superdex200-increase 10/300 GL SEC column (GE Healthcare) pre-equilibrated with nanodisc buffer: 25 mM Tris, pH 7.4, 150 mM NaCl. Fractions containing PhuR nanodiscs were pooled, concentrated, flash frozen in LN2, and stored at -80°C until further use.

### Identification of PhuR-specific sABs using phage display

For sAB screening, biotinylated nanodiscs containing PhuR were used. Selections were performed using PhuR both from native and refolding conditions. Prior to phage display, the efficiency of capture on streptavidin-coated paramagnetic beads (Promega) was evaluated as described before. Biopanning was performed using phage display sAB Library E (kind gift of S. Koide) in selection buffer (25 mM Tris, pH 7.4, 150 mM NaCl, 1% BSA) according to previously described protocol^17–19^. In the first round, biopanning was performed manually using 400 nM of PhuR nanodiscs immobilized onto paramagnetic beads by one-hour incubation at RT with Library E. Following three washes with selection buffer, beads enriched for phage displaying PhuR-specific sABs were used to directly infect log-phase *E. coli* XL-1 Blue bacteria. Following a 20-min infection, 2×YT media supplemented with ampicillin (100 µg/mL) and M13-KO7 helper phage (10^9^ PFU/mL) was added and phage were amplified overnight. To increase the stringency of sAB selection, four additional rounds of biopanning were performed by decreasing PhuR concentrations in a stepwise manner: 200, 100, 50, and 25 nM. For rounds 2-5, the amplified phage pool from each preceding round was using as the input, and biopanning was performed semi-automatically using a Kingfisher magnetic beads handler (Thermo Fisher Scientific). The phage pools for rounds 2-5 were pre-cleared with 200 µL of streptavidin beads to remove non-specific binders. In all rounds, a tenfold molar excess of non-biotinylated E3D1 nanodiscs were used as soluble competitors to remove MSP-specific sABs. Finally, for rounds 2-5 phage particles were eluted from magnetic beads by a 15-min incubation step with 1% Fos-choline-12 (Anatrace) prepared in buffered saline.

### Initial validation using single-point phage ELISA

Colonies of E. coli XL-1 Blue bacteria containing individual phagemid clones from round 5 of phage display were used to inoculate 400 µL of 2×YT media supplemented with ampicillin (100 µg/mL) and M13-KO7 helper phage (10^9^ PFU/mL). Phage were amplified overnight in 96-well deep well blocks at 37°C with shaking at 280 RPM. Amplified phage were diluted into ELISA buffer (selection buffer with 0.5% BSA) and then tested against PhuR-loaded or empty biotinylated E3D1 nanodiscs. All ELISA experiments were performed on 96-well ELISA plates (Nunc) coated overnight with 2 µg/mL neutravidin and blocked for at least two hours with selection buffer. ELISA were performed as previously described and bound phage were detected with TMB substrate (Thermo Fisher Scientific) following a 30-min incubation with HRP-conjugated anti-M13 monoclonal antibody (Antibody Design Laboratories)^19^. Absorbance was measured at 450 nm following quenching with 1 M HCl. Wells containing empty biotinylated E3D1 nanodiscs were used to determine non-specific binding. Specific binders were determined based on their signal/background ratio.

### sAB expression and purification

Selected clones based on phage ELISA were sequenced at the University of Chicago Comprehensive Cancer Center DNA Sequencing facility. Unique clones were sub-cloned into the sAB expression vector pRH2.2 (kind gift of S. Sidhu) using the In-Fusion Cloning Kit (Takara Bio). The best candidates (based on highest ELISA signals) were expressed and purified as previously described^19,51^.

### Assessment of sAB binding affinity for PhuR

ELISA were performed using purified sABs to estimate apparent binding affinities as previously described. ELISA plates were prepared as above for phage ELISA. 25 nM of either PhuR-loaded or empty biotinylated nanodiscs were immobilized onto wells. Binding was assessed using 3-fold serial dilutions of sABs starting from 3 µM maximum concentrations. Bound sABs were detected with TMB substrate following a 30-min incubation with HRP-conjugated recombinant protein L (Pierce). Reactions were quenched with 1 M HCl and absorbance was measured at 450 nm. The signals were background subtracted and normalized A_450_ values were plotted against the log_10_ sAB concentrations. EC_50_ values were then calculated in Prism 9 (GraphPad) using a variable slope model and assuming a sigmoidal dose response.

### Amphipol (APol) reconstitution

APol A8-35 (Anatrace) was mixed with native or refolded PhuR in SEC buffer with DDM in a 1:10 PhuR/APol mass ratio. The solution was incubated on ice for 20 minutes, followed by the addition of Bio-Beads to remove DDM. Amphipol reconstitution was performed overnight at 4°C with nutation. The reconstitution mixture was then transferred from Bio-Beads, filtered using 0.2-µm centrifugal filter units, and injected onto a Superdex200-increase 10/300 GL column equilibrated with SEC buffer (without DDM). Fractions containing PhuR reconstituted into A8-35 were analyzed by SDS-PAGE, pooled, concentrated, and flash frozen until further use. An identical protocol, except using biotinylated A8-35 (Anatrace), was used to reconstitute refolded PhuR into biotinylated amphipols for phage display.

### Heme binding studies

5-µM of protein solution in APols was prepared and added to a 96-well plate. Hemin (Sigma) was dissolved in 0.1 M NaOH and shaken at 1000 RPM for 10 min. The resulting suspension was then centrifuged (16000×g, 10min) and passed through a 0.22 μm filter. The concentration was determined by measuring absorbance (385 nm) of 1000X diluted solution and the heme molar extinction coefficient (ε385 = 58440 M-1 × cm-1). Hemin solutions ranging from 1-50 µM were added to PhuR and incubated for 20 minutes at RT. The UV-Vis spectrum was recorded from 190-850 nm using a Nanodrop One (Thermo Fisher Scientific). The background signal was subtracted from protein-heme wells using wells containing heme but not protein. Values at the Soret peak (410 nm) were plotted as a function of heme concentration and analyzed using nonlinear regression in Prism 9 (GraphPad). Additionally, the spectrum was recorded for the complex of native PhuR, sAB11, and nanobody that was used to prepare the cryo-EM sample (∼5μM total protein concentration). These data were used for comparison with spectra collected from the heme binding experiments.

### PhuR complex formation with sAB11 and anti-sAB nanobody

Amphipol-reconstituted native PhuR (10 µM) was combined with a 2-fold molar excess of sAB11 and 3-fold molar excess of an anti-sAB nanobody, which was expressed and purified as previously described. Following a 20 minute incubation, the complex was loaded on a Superdex200-increase 10/300 GL column (Cytiva) in SEC buffer (without DDM). Fractions containing all complex components were pooled and concentrated resulting in A_280_ 6.9, which was determined to be is 4.2 mg/mL based on the theoretical extinction coefficients of the individual components.

Complexes with amphipol-reconstituted refolded PhuR with sAB11 were similarly formed and isolated by using 1 µM of refolded PhuR, except for the addition of 50µM hemin prior to incubation with the sAB and nanobody.

### Negative-stain electron microscopy

To determine sample quality and success of complex formation, purified proteins were diluted to 0.05 mg/mL and applied onto plasma-cleaned (Gatan Solarus) copper grids with a continuous carbon layer (Electron Microscopy Sciences). The grids were washed twice with droplets of water, and then blotted with Whatman 4 filter paper to remove the excess solution. The grids were stained with two applications of 1% uranyl formate (EMS), and again blotted. After drying, the grids were imaged using a FEI Spirit 120 kV microscope equipped with a digital CCD camera with 2k × 2k resolution using a pixel size of 2.1 Å. In total, 30-50 images were collected for the native and refolded PhuR with and without sABs, using peak SEC fractions diluted into SEC buffer at the aforementioned concentration to prepare the ns-EM grids. Images were processed using Relion-3.1 to obtain two-dimensional class averages^52^.

### Single-particle cryo-EM vitrification and data collection

APol-reconstituted native PhuR complexes with sAB11 and the nanobody were concentrated to 1.1-4.2 mg/mL using a 100-kDa molecular weight cutoff concentration (Amicon). Prior to vitrification, 0.25% CHAPSO was added to improve the quality of vitreous ice during freezing and reduce denaturation and aggregation at the air-water interface. 3.5 µL of sample was applied to plasma-cleaned (Gatan Solarus) 1.2/1.3-µm UltrAuFoil grids (Quantifoil) and blotted using standard Vitrobot filter paper (Ted Pella, 47000-100) on both sides for 4 seconds using a Vitrobot Mark IV (Thermo Fisher Scientific) operating at 8°C and 100% humidity. The grids were immediately plunged after blotting into liquid ethane for vitrification. Cryo-EM images were collected at the University of Chicago Advanced Electron Microscopy Facility using a Titan Krios G3 electron microscope (Thermo Fisher Scientific) operating at 300 kV and equipped with a Gatan K3 direct detection camera and Energy Filter (BioQuantum). Automated data collection was done in CDS mode using the EPU software package. A total of 5,031 movies consisting of 50 fractions with a 60 e^-^/Å total exposure were collected with a super-resolution pixel size of 0.534 Å, a calibrated pixel size of 1.068 Å, and a defocus range of -1.0 to -2.5 µm.

### Cryo-EM data processing for PhuR-sAB11-nanobody complex

All data processing and visualization software used were part of the SBGrid collection except for cryoSPARC (v3.3)^53,54^. Movies were subjected to motion correction by MotionCor2 (v1.4.0) within the Relion-4.0-beta GUI^55–57^. CTF parameters were determined using GCTF (v1.06)^58^. Micrographs were filtered based on defocus values, astigmatism, resolution, and figure of merit, leaving 4,153 micrographs in the final set. Around 5 million particles were selected using the Laplacian-of-Gaussian (LoG) picker. The particles were extracted using a box size of 360 pixels and imported into cryoSPARC (v3.3). Several iterations of 2D class averaging were performed to remove junk particle. Particles were selected based on whether 2D classes contained monomers or antiparallel dimers of PhuR. These sets were independently and in parallel subjected to *Ab initio* reconstruction, classification using heterogeneous refinement, and non-uniform refinement, with the final particle sets containing 277,088 and 141,268 particles and producing monomer and dimer maps of 3.0 and 3.3 Å resolutions, respectively. The particle sets were exported to Relion for CTF refinement and Bayesian polishing using pyem, and then imported back to cryoSPARC for non-uniform refinement and local refinement using masks generated in Chimera around PhuR and the sAB variable domains^59,60^. This resulted in maps with 2.9 and 3.0 Å for the monomer and dimer, respectively.

To obtain a higher number of “good” particles, template-based picking was performed in Relion by first generating 2D classes using the monomer particle stack. 2.4 million particles were selected and subjected to rounds of 2D class averaging and 3D refinement as above. Following CTF refinement and Bayesian polishing in Relion of the best class of PhuR monomers, non-uniform refinement in cryoSPARC produced a 2.6 Å map. This map was subjected to local refinement in cryoSPARC using a tight mask around PhuR and the sAB variable region prepared in Chimera (v1.15), which produced a 2.5 Å map. Local resolution maps were prepared in cryoSPARC and visualized using ChimeraX (v1.5)^61,62^. The detailed data processing workflow is shown in **Supplemental Fig. 3**. Resolutions are reported based on the Fourier Shell Correlation (FSC) equals 0.143 criterion.

### Structural model building and refinement

The *B*-factor sharpened map from the cryoSPARC local refinement job was used for model building of the PhuR-sAB11 complex. The AlphaFold model of PhuR was used as the starting coordinates, and manually adjusted into the cryo-EM density map using Chimera. The manually adjusted model and map were then brought into Coot (v0.9) and further adjusted into the density using morphing^63^. The sAB variable region was built using a template (5BJZ) with the CDRs removed, and the sAB11 CDRs were manually built in Coot using the sequence and cryo-EM density to guide manual fitting. PhuR residues without experimental cryo-EM density were deleted. Heme and the lipopolysaccharide (LPS) were built in Coot by manual adjustment into the density. Coordinates for heme were obtained from the Refmac ligand library, and LPS coordinates were obtained from PubChem^64,65^. The LPS headgroup was truncated to the first KDO sugar due to the absence of cryo-EM density for this region. The combined PhuR-sAB model with bound ligands was subjected to multiple rounds of refinement in Coot and Phenix (v1.20), with the map sharpened or blurred in Coot to facilitate model building^66^. The model was inspected manually after each cycle, and refinement was performed until no further improvement was observed. Waters were added into non-protein and non-ligand densities using Douse and manually inspected. The models were validated in Phenix using its comprehensive cryo-EM validation tools, MolProbity, and EMRinger^67,68^. Directional anisotropy of the map was evaluated using 3DFSC^69^.

### Structural Analyses

Protein surface interactions were calculated using the Proteins, Interfaces, Structures, and Assemblies server and visualized using PyMol^70^. Consurf was used to analyze evolutionary conservation using the PhuR structure^45,46^. Cavities were analyzed using the Voss Volume Voxelator online tool (http://3vee.molmovdb.org/)^71^. Heme interactions were visualized using LigPlot+^72^. All other images were prepared using Chimera and ChimeraX.

## QUANTIFICATION AND STATISTICAL ANALYSIS

- Cryo-EM data collection and refinement statistics are presented in Table 1.
- Heme binding assays, performed in triplicate, are expressed as mean ± SEM. Experiment details are in Methods.
- Supplementary Table 2 includes estimated sAB affinities from ELISA and non-linear regression analysis. Refer to Methods for experimental details.

