## Supplemental Material for "Chaperone-assisted cryo-EM structure of *P. aeruginosa* PhuR reveals molecular basis for heme binding"

**Table S1.** CDR sequences of sABs for selected clones. Sequences of CDRs diversified in Library E shown here[1]. Number of times that the clone appeared in the final phage pool is shown as repeat number; related to Figure 1

| sAB ID | L3 aa | HC1 aa | HC2 aa | HC3 aa | Repeat No. |
|---|---|---|---|---|---|
| sAB1 | SSSSLI | FSSSYI | SISPYYGYTS | EEYSYWWKWFISHYGL | 1 |
| sAB2 | SSWSTPI | IYSYSI | SIYPSYSSTS | GEYMWGYWQWYYYDEWAF | 1 |
| sAB3 | SEQFYPI | IYSSSI | SISPYSGYTS | GTLYWAMFWGGGGYWGL | 1 |
| sAB4 | YLSRSLI | FSYSSI | SISSSSGSTS | GYHSIWRYGYAF | 1 |
| sAB5 | SSSSLI | VSYYYI | SISPYSGSTS | GYWDAWYYMWSYPQSWEGM | 1 |
| sAB6 | SSSSLI | FSSSSI | SISSSSGSTS | HYYDPWWFWYMAHAM | 1 |
| sAB7 | YLWEQLV | VYYSSI | SISSYYGSTS | MGEYMGWPYAF | 2 |
| sAB8 | YYFWYPI | IYSYYI | SIYSYSGSTS | QVQYYAWSWYTEKWGAAM | 1 |
| sAB9 | SSSSLI | VSYYSI | SISSSSGSTS | RGLDWWWYWSYSYAYGL | 4 |
| sAB10 | FVYGQLI | VSYSSI | YISSYSGYTY | RSGWAM | 1 |
| sAB11 | GSSSPL | FYYYSI | SISPYYGSTY | SMNWYYSSPYGMSQGM | 2 |
| sAB12 | SFFSGLI | VSYSSI | SIYSSSGYTS | SVSWSWYWLSAF | 1 |
| sAB13 | SSSSLI | FSSSSI | SISSSSGSTS | SYLYSWYHFWAWYLSGGF | 2 |
| sAB14 | SWGGSLL | FSSSSI | SISSSSGSTS | YKKYAWYMSWAYPSAI | 2 |
| sAB15 | SSSSLI | FSSSSI | SISSSSGSTS | YKYYSYMWFVGIYSHAL | 5 |
| sAB16 | SSSSL | FSSSSI | SISSSSGSTS | YSAYSWFFPMSAM | 3 |
| sAB17 | SSSSLI | FSSSYI | SISSSSGSTS | YSYSSWWFPMKAL | 5 |
| sAB18 | SSSSLI | FSSSSI | SISSSSGSTS | YYSAWWYYWHDYGYWYGM | 5 |

**Table S2.** Table of $EC_{50}$ and $R^2$ values obtained from affinity-estimation ELISA for selected sAB clones; related to Figure 1

| sAB | 8 | 3 | 5 | 11 | 2 | 6 |
|---|---|---|---|---|---|---|
| EC50 | 2.21E-08 | 2.22E-08 | 1.84E-08 | 8.75E-09 | 1.20E-08 | 2.55E-07 |
| $R^2$ value | 0.9101 | 0.9868 | 0.9328 | 0.9894 | 0.9733 | 0.9603 |

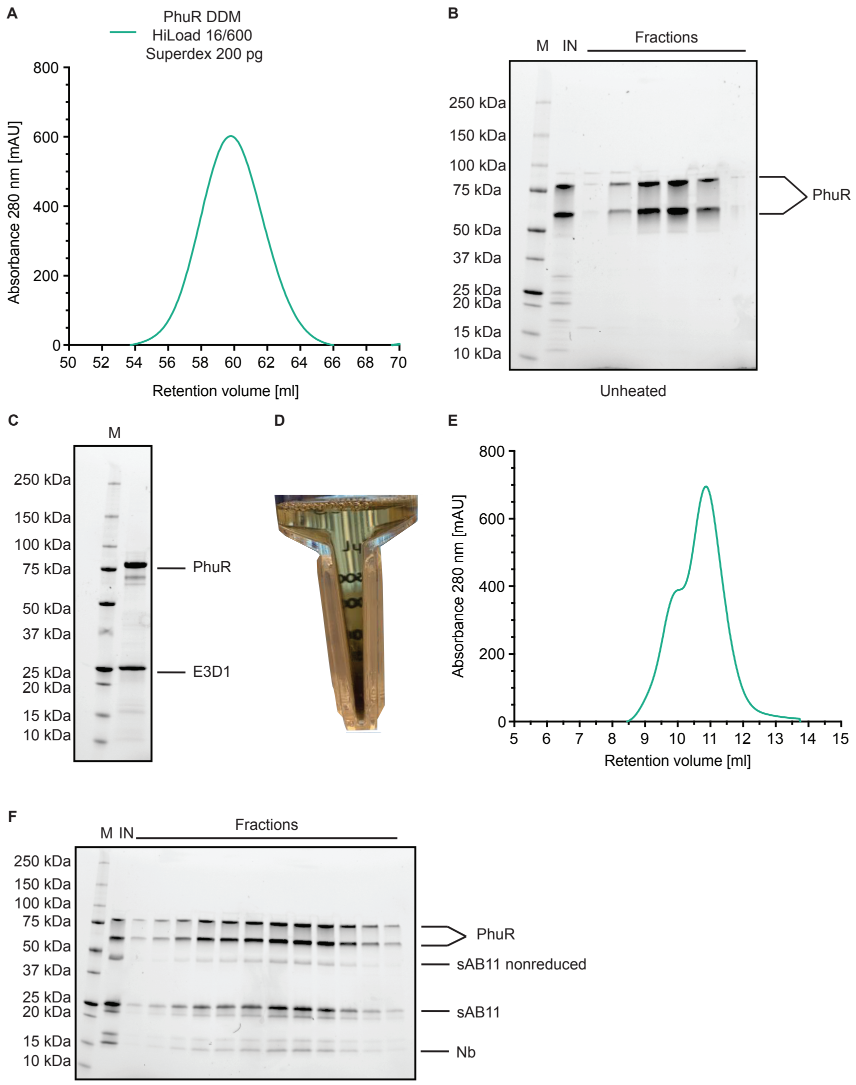


**Figure S1**. PhuR purification in DDM, nanodisc reconstitution and cryoEM sample preparation; related to Figure 1, 2, 3, 4. **A.** SEC profile of DDM solubilized protein using pooled PhuR-containing fractions from initial IMAC purification. **B.** SDS-PAGE gel showing protein-containing fractions from SEC run shown in panel A. PhuR appears as two bands due to the heat-modifiability property of β-barrel outer membrane proteins[2]. M - marker; IN - input followed by fraction number corresponding to panel A X-axis. **C.** SDS-PAGE gel of PhuR reconstituted into E3D1 nanodiscs; M – marker. **D.** Amicon centrifugal filter containing concentrated DDM solubilized PhuR solution. The brown solution color comes from the presence of heme. **E.** SEC profile of amphipol solubilized PhuR in complex with sAB11 and nanobody. **F.** SDS-PAGE gel showing protein containing fractions from SEC run shown in panel E. M – marker; IN – input followed by collected fractions. The heavy and light chains of sAB11 run as two closely spaced bands around ~24-25 kDa, with a small population of non-reduced sAB at ~50 kDa. The Nb similarly runs as two bands due to a mixture of reduced and non-reduced species.

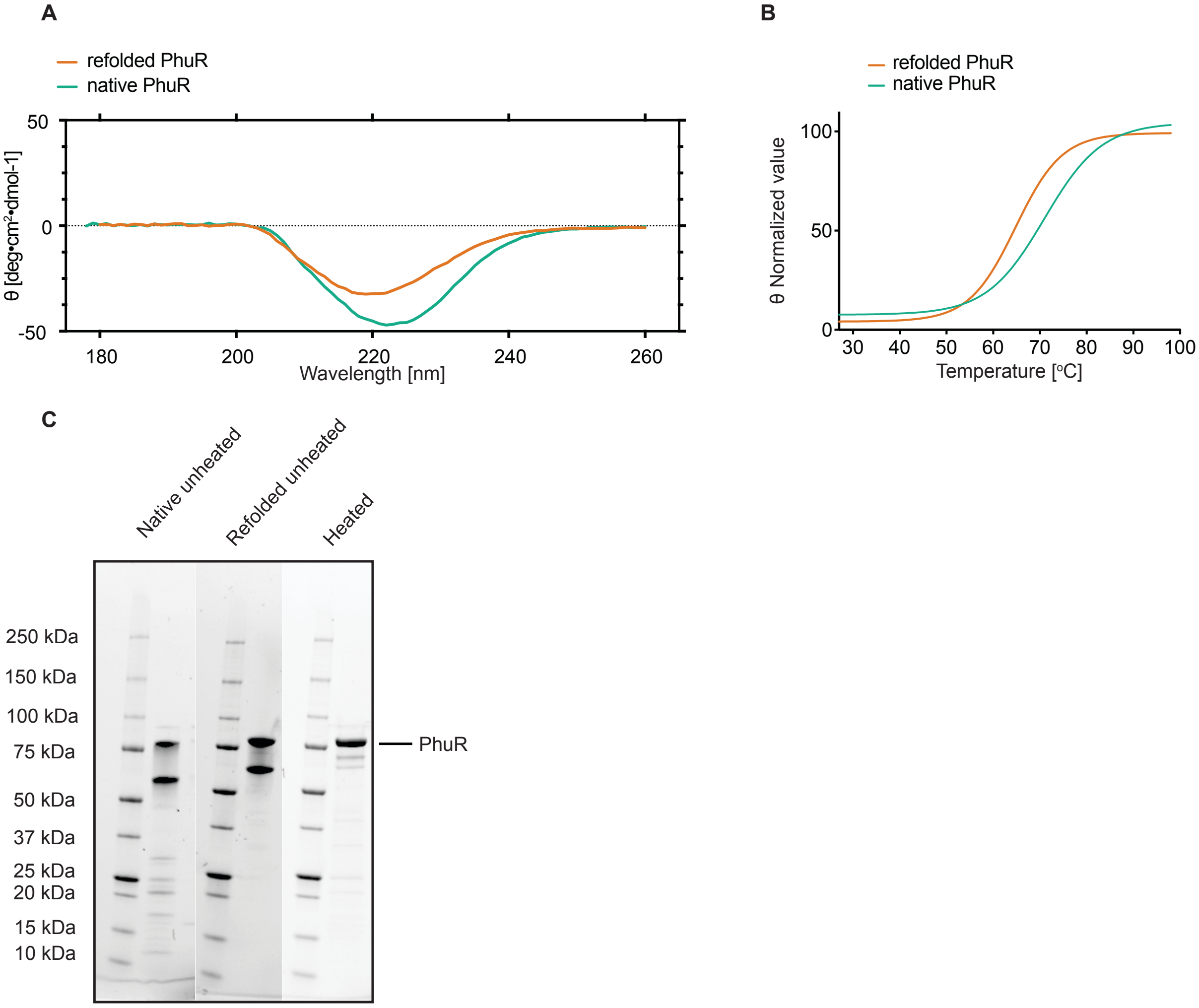


**Figure S2.** Secondary structure and thermal stability of native and refolded PhuR; related to Figure 5. **A.** Far-UV CD spectra of refolded (orange) and native (green) PhuR. Graphs plot molar ellipticity vs. wavelength. **B.** Normalized molar ellipticity at 222 nm plotted as a function of temperature to show thermal stability of native (green) and refolded PhuR (orange). This wavelength was monitored due to its signature for β-barrel proteins. Curves fit using Prism 9.0 to determine melting temperature. Thermal denaturation experiments of refolded and native PhuR showed melting temperature of 65°C and 71°C respectively. **C.** SDS-PAGE gel showing unheated refolded and native DDM solubilized PhuR samples and heated native PhuR. Unheated samples run as two bands due to the heat modifiability property of β-barrel outer membrane proteins. The proportion of higher molecular weight band, corresponding to the fully denatured PhuR, is higher in the refolded sample than the native, suggesting a greater proportion of the sample is more readily denatured and hence less stable in the former condition. The image is a composite of multiple gels, which were cropped and assembled to highlight the most relevant lanes and bands.

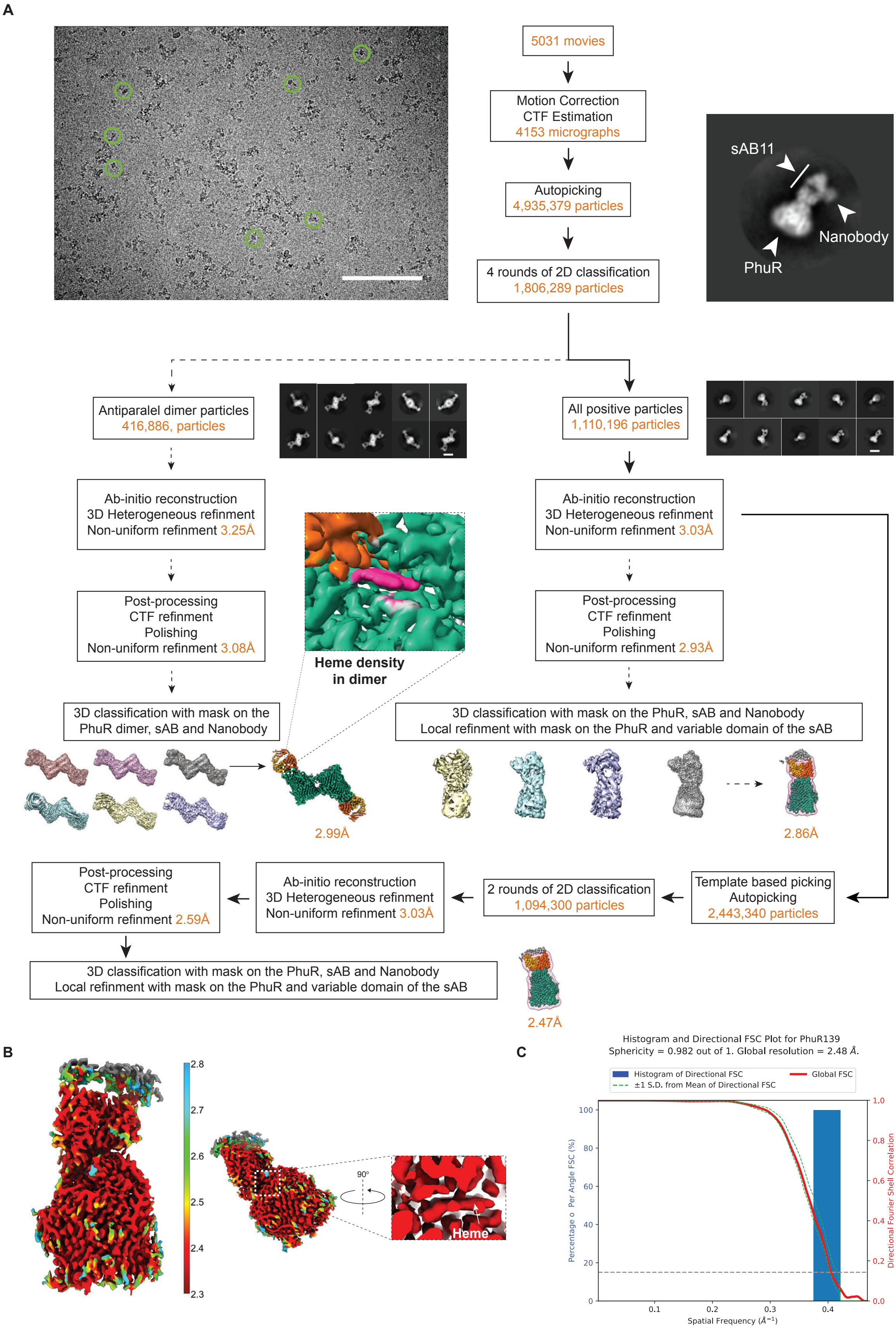
A
5031 movies
Motion Correction
CTF Estimation
4153 micrographs
Autopicking
4,935,379 particles
4 rounds of 2D classification
1,806,289 particles
sAB11
Nanobody
PhuR
Antiparalel dimer particles
416,886, particles
All positive particles
1,110,196 particles
Ab-initio reconstruction
3D Heterogeneous refinment
Non-uniform refinment 3.25Å
Ab-initio reconstruction
3D Heterogeneous refinment
Non-uniform refinment 3.03Å
Post-processing
CTF refinment
Polishing
Non-uniform refinment 3.08Å
Heme density
in dimer
Post-processing
CTF refinment
Polishing
Non-uniform refinment 2.93Å
3D classification with mask on the
PhuR dimer, sAB and Nanobody
3D classification with mask on the PhuR, sAB and Nanobody
Local refinment with mask on the PhuR and variable domain of the sAB
2.99Å
2.86Å
Post-processing
CTF refinment
Polishing
Non-uniform refinment 2.59Å
Ab-initio reconstruction
3D Heterogeneous refinment
Non-uniform refinment 3.03Å
2 rounds of 2D classification
1,094,300 particles
Template based picking
Autopicking
2,443,340 particles
3D classification with mask on the PhuR, sAB and Nanobody
Local refinment with mask on the PhuR and variable domain of the sAB
2.47Å
B
2.8
2.7
2.6
2.5
2.4
2.3
90°
Heme
C
Histogram and Directional FSC Plot for PhuR139
Sphericity = 0.982 out of 1. Global resolution = 2.48 Å.
Histogram of Directional FSC
±1 S.D. from Mean of Directional FSC
Global FSC
Percentage o Per Angle FSC (%)
Directional Fourier Shell Correlation
Spatial Frequency (Å⁻¹)
0
20
40
60
80
100
0.0
0.2
0.4
0.6
0.8
1.0
0.1
0.2
0.3
0.4

**Figure S3.** Cryo-EM single particle analysis of amphipol-reconstituted PhuR-sAB11-Nb complex; related to Figure 2. **A.** Cryo-EM 3D reconstruction workflow. Representative micrograph shown with 50 nm scale bar and selected particles are shown in green circles. Select 2D class averages shown for antiparallel dimer of PhuR (left) with close-up view of density for bound heme and monomeric PhuR (right) with indicated sAB and Nanobody (top). Arrows show progress of reconstruction. Dashed lines show initial data processing strategy. Solid lines show final reconstruction strategy. **B.** Cryo-EM density map of PhuR colored by local resolution with close-up view of density for bound heme. **C.** Directional Fourier Shell Correlation (FSC) plot showing global FSC (red line) with histogram of directional FSCs (blue bars) and standard deviation from mean FSC[3]. Global resolution reported using the FSC = 0.143 criterion.

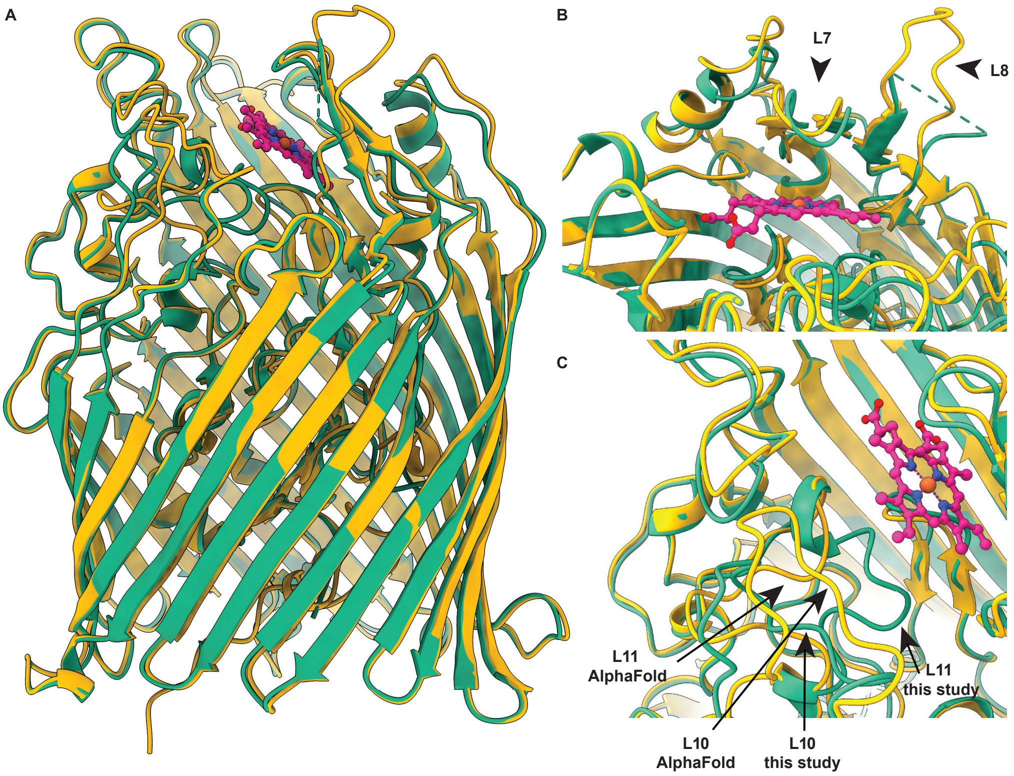


**Figure S4:** Comparison of PhuR structure (this study) and AlphaFold PhuR model[4]; related to Figure 2, 3. **A.** Side view of aligned experimental PhuR structure (green) and AlphaFold PhuR model (yellow), with heme molecule in magenta represented as sticks and balls. **B.** Close up view of conserved FRAP/PNPNL and unresolved loops (L7 and L8) located above the bound heme molecule. **C.** Close up view of L10 and L11 loop to highlight conformational differences between the experimental structure and model. In the cryoEM structure, L10 is oriented towards the vestibule, enclosing the heme ligand in its binding pocket. Black arrows highlight the differences between the experimental structure and model.

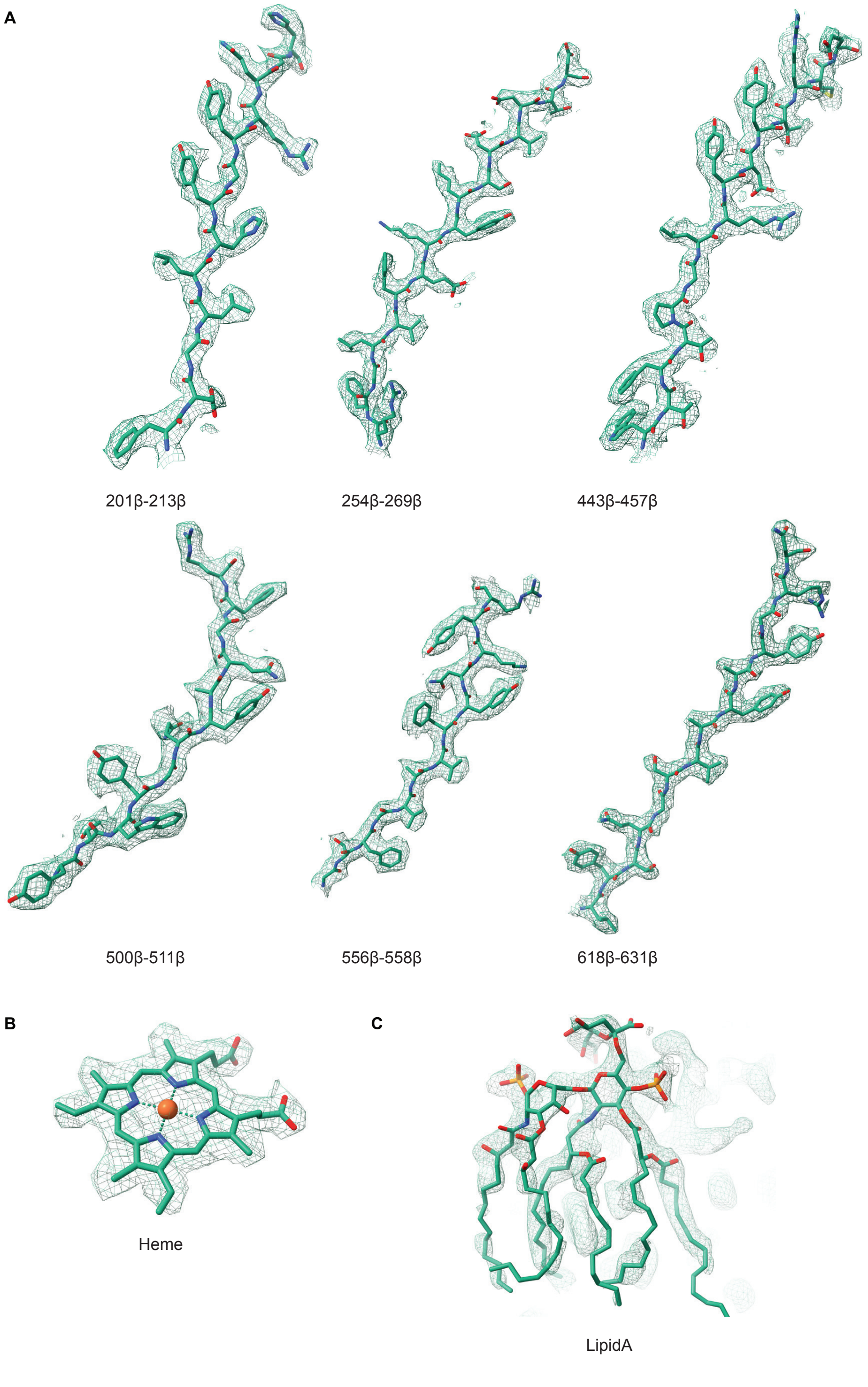


**Figure S5.** Model in map fit of PhuR-sAB11-Nb complex; related to Figure 2, 3. **A.** Selected β-strands from barrel domain shown as green sticks overlaid with cryo-EM density (green mesh). **B.** Heme ligand overlaid with cryo-EM density and represented as in A. **C.** LPS represented as green sticks and overlaid with cryo-EM and colored as in A.

A

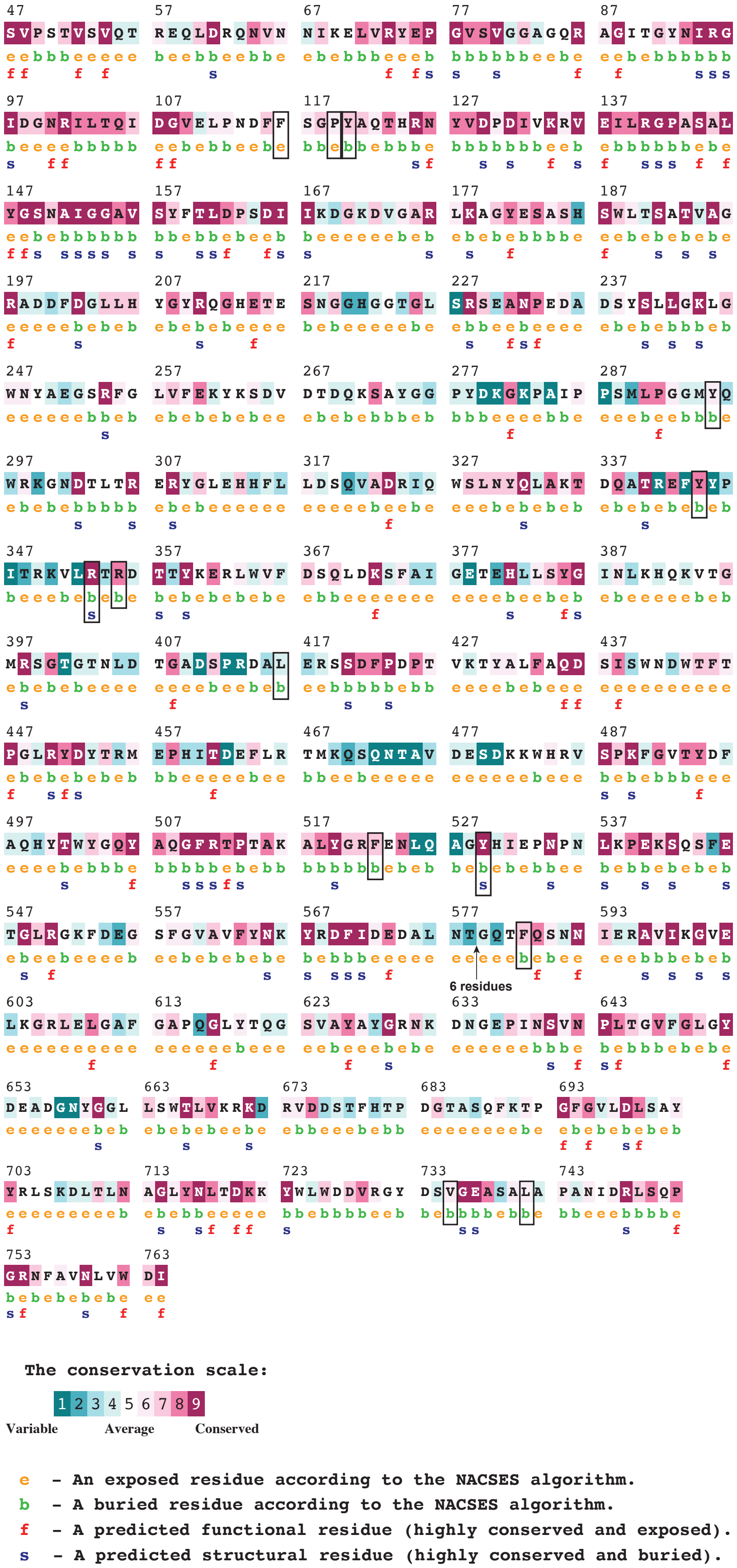
The conservation scale:
1 2 3 4 5 6 7 8 9
Variable
Average
Conserved
e - An exposed residue according to the NACSES algorithm.
b - A buried residue according to the NACSES algorithm.
f - A predicted functional residue (highly conserved and exposed).
s - A predicted structural residue (highly conserved and buried).

**Figure S6.** ConSurf evolutionary analysis of PhuR; related to Figure 2, 3. **A.** Individual residues from cryo-EM structure of PhuR colored by conservation score. Residues involved in heme interactions are highlighted with boxes. Residues that were not modeled in the structure were excluded by the software from the analysis and are indicated between T578 and G585 with the rest numbered as in the structure. Residues labeled by predicted surface exposure and functionality as based on ConSurf analysis[5,6].

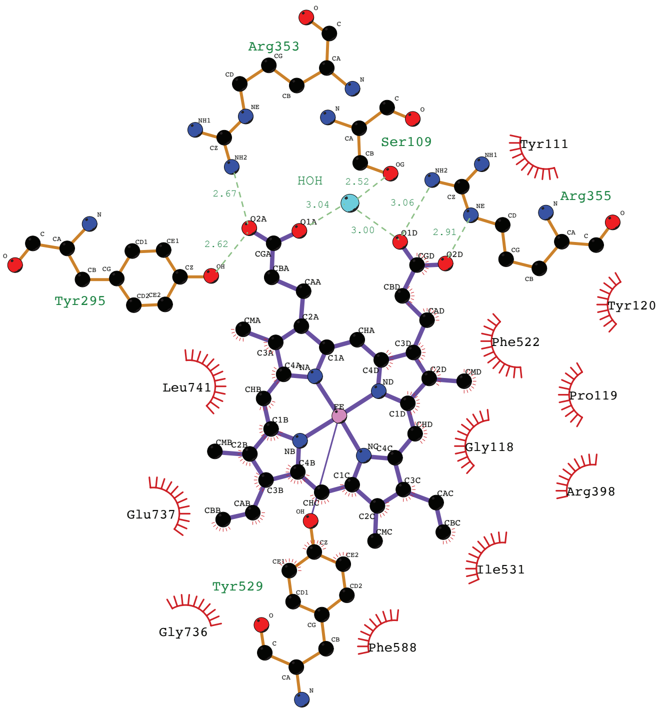


**Figure S7.** Residue interactions with heme within the extracellular vestibule as based on LigPlot[7]; related to Figure 3.
